# Time-course transcriptomic dataset of gallic acid-induced human cervical carcinoma HeLa cell death

**DOI:** 10.1101/2025.02.03.636357

**Authors:** Ho Man Tang, Peter Chi Keung Cheung

## Abstract

Gallic acid is a natural phenolic acid that displays potent anti-cancer activity in clinically relevant cell culture and rodent models. Although research has focused on determining the efficacy of gallic acid against various types of human cancer cells, the molecular mechanisms governing the anti-cancer properties of gallic acid remains largely unclear, and a transcriptomic study of gallic acid-induced cancer cell death has rarely been reported. Therefore, we performed time-course bulk RNA-sequencing study to elucidate the molecular signature of gallic acid-induced cell death in human cervical cancer HeLa cells, as this is a widely used *in vitro* model in the field. Our RNA-sequencing dataset covers the early (2^nd^ hour), middle (4^th^, 6^th^ hour), and late (9^th^ hour) stages of the cell death process after exposure of HeLa cells to gallic acid, and the untreated (0^th^ hour) cells served as controls. Differential expression of messenger RNAs (mRNA) and long non-coding RNAs (lncRNA) were identified in each time point in the dataset. In summary, this dataset is a unique and valuable resource with which the scientific community can explore the molecular mechanisms and identify key regulators of the gallic acid-induced cancer cell death process.

## Background & Summary

Gallic acid (GA), also known as 3,4,5-trihydroxybenzoic acid, is a natural bioactive substance that displays anti-cancer properties^1-4^. This low molecular weight phenolic compound (C7H6O5) is commonly found in edible fungi, fruits, nuts, vegetables, tea leaves, and herbs^5,6^. Accumulating evidence from cell culture and clinically relevant rodent models has demonstrated the anti-cancer bioactivity of gallic acid, and its derivatives, in various human cancer cell types^1-4^, such as adenocarcinoma^7^, bone cancer^8^, breast cancer^9^, cervical cancer^10^, colon cancer^11^, gastric cancer^12^, leukemia^13^, liver cancer^11^, lung cancer^14^, lymphoma^15^, ovarian cancer^16^, prostate cancer^17^, and skin cancer^18^. Importantly, gallic acid is well-tolerated by rodent models at the dosages that display anti-cancer effect^19-23^. Thus, both *in vitro* and *in vivo* studies reveal the pharmacological potential of developing gallic acid as a broad-spectrum anti-cancer agent for humans, one that is expected to be safely applied to treat cancers, with minimal side effects.

Resisting cell death is the hallmark of cancer^24,25^. To overcome the obstacle in drug resistance, the ideal anti-cancer agents should therefore be able to simultaneously activate multiple pathways that trigger cancer cell death^26,27^. The three major targets of cancer therapy are a cell suicide pathway called apoptosis^28^, an iron-dependent form of non-apoptotic cell death called ferroptosis^29^, and a regulated form of necrotic cell death called necroptosis^30^. Interestingly, we have recently shown that gallic acid can trigger concurrent apoptosis, ferroptosis, and necroptosis in human cancer cell lines^31,32^. Our finding provides new mechanistic insights into the high potency of gallic acid in killing cancer cells reported by other investigators in a diversity of *in vitro* and *in vivo* models^1-4^. As apoptosis, ferroptosis, and necroptosis regulate cell death by distinctive mechanisms^33-35^, the cancer cells that resist one of those cell death mechanisms can be still killed by gallic acid because it activates all three of these cell death pathways.

To unleash the potential of gallic acid in clinical application, it is important to elucidate the molecular mechanisms that govern gallic acid-induced cancer cell death. Therefore, we performed a time-course RNA-sequencing (RNA-Seq) to reveal the transcriptomic profile of the early (2^nd^ hour), middle (4^th^, 6^th^ hour), and late (9^th^ hour) stages of gallic acid-induced cell death process^32^. We used human cervical cancer HeLa cells as the study model here, because this cell line is commonly used in studies of gallic acid and cancer biology^1,36^. We applied 50μg/mL of gallic acid to trigger HeLa cell death in this study, as this is the physiologically relevant dosage that is lower than the no-observed-adverse-effect level (NOAEL) in the rodent models used for studying gallic acid^19-21^, and this dosage displays anti-cancer activity to a wide range of cancer cell types *in vivo* and *in vitro*^1-4^.

Of particular interest, our RNA-Seq dataset^32^ covers both the well characterized protein-coding RNAs/ messenger RNAs (mRNA)^37^, and the long non-coding RNAs (lncRNA)^38^ in which their functions are being emerged. Therefore, our dataset will serve as a unique and valuable resource for further studying the molecular mechanisms that regulate the anti-cancer activity of gallic acid, thereby facilitating the translation of this natural anti-cancer agent into clinical applications.

### Methods

#### Cell culture

The human cervical cancer HeLa cell line was obtained from the American Type Culture Collection (ATCC, Manassas, VA, USA; catalog # CCL-2). This cell line was cultured in Dulbecco’s Modified Eagle Medium/Nutrient Mixture F-12 (DMEM/F-12; catalog # 10565042) supplemented with 10% heat inactivated fetal bovine serum (FBS; catalog # 16140071), 2mM GlutaMAX (catalog # 35050061), 100 U/mL penicillin and 100 μg/mL streptomycin (catalog # 15070063) purchased from Thermo Fisher Scientific (Waltham, MA, USA), at 37 degree Celsius (°C) under an atmosphere of 5% carbon dioxide (CO2). Before the start of the experiment, HeLa cells were plated onto a 100 mm Corning tissue culture dish (Corning, NY, USA; catalog # 430167) for 1 day to reach 80% cell confluency.

#### Cell death induction by gallic acid

Gallic acid (catalog # G7384) dissolved in dimethyl sulfoxide (DMSO; catalog # D2438) purchased from Sigma-Aldrich (Burlington, MA, USA) was to make a 20 mg/mL stock solution. This was then further diluted into the cell culture medium (described above) to achieve the working concentration of 50 μg/mL, before being applied to the cells to induce cell death. The stock solution was freshly made before each cell death induction experiment.

#### Total RNA isolation

Total RNA was isolated from the HeLa cells that were exposed to 50 μg/mL gallic acid for 0, 2, 4, 6, and 9 hours (**Fig. 1**). At each of the timepoints, the cell culture medium was removed from the cell culture dish, and the cells were lysed on the dish using 2 mL of ice-cooled TRIzol by thorough pipetting. Total RNA was extracted using the Direct-zol™ RNA MiniPrep kit (Zymo Research, Irvine, CA, USA; catalog # R2070T) according to the manufacturer’s instructions, eluted in nuclease free water, quantified by a NanoDrop ND-1000 spectrophotometer (NanoDrop Technologies, Wilmington, DE, USA), and stored at -80°C.

**Figure 1.**
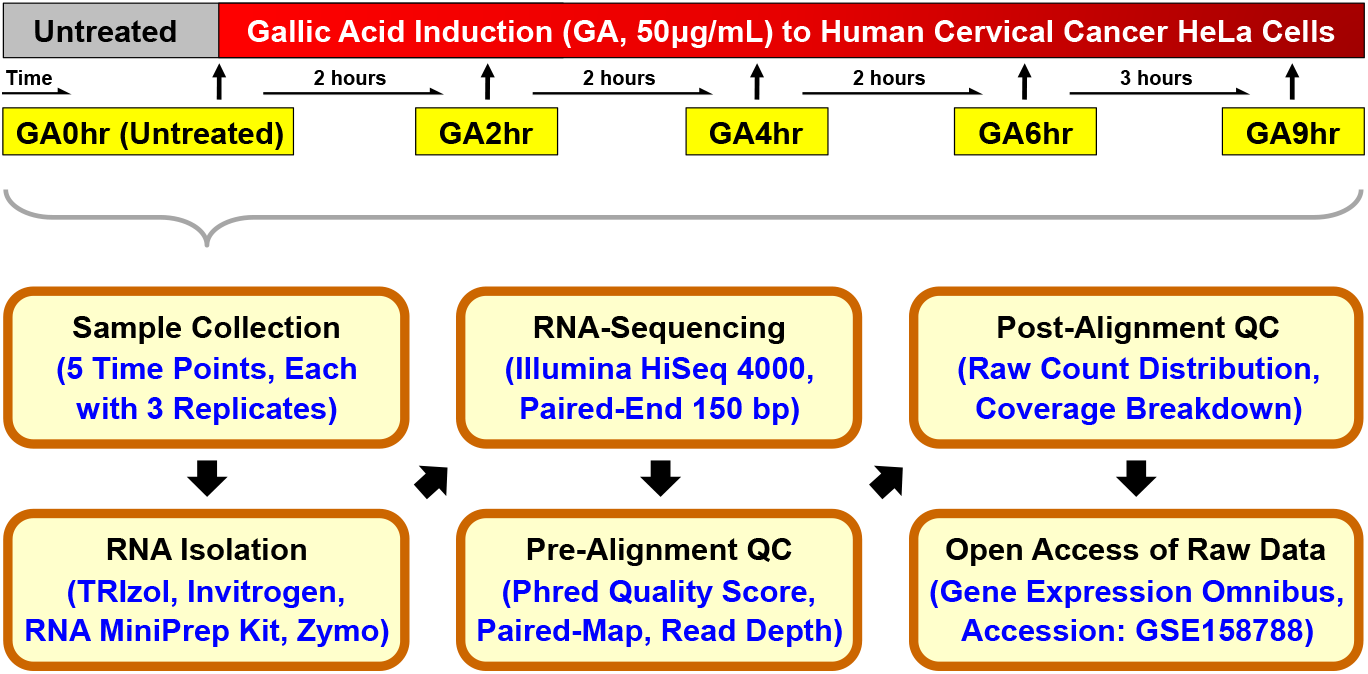
Schematic of the workflow for the time-course RNA-sequencing study. Human cervical cancer HeLa cells were treated with 50μg/mL gallic acid for 2 hours (GA2hr), 4 hours (GA4hr), 6 hours (GA6hr), and 9 hours (GA9hr) to induce the cell death process. Untreated cells served as control (GA0hr). Cell samples of three biological replicates from each condition mentioned above were collected for total RNA extraction, analysed using Illumina HiSeq 4000 for RNA-sequencing, and subjected to pre-alignment and post-alignment quality checks. The raw RNA-sequencing dataset of all 15 samples is publicly available at the Gene Expression Omnibus (GEO accession: GSE158788)^49^.

#### RNA-sequencing

The RNA-sequencing analysis was conducted at the Novogene Core Facility (Novogene Corporation Inc., Sacramento, CA, USA) using the standard protocols from Novogene and Illumina (San Diego, CA, USA)^32^. The RNA concentration was measured by using a NanoDrop spectrophotometer and Qubit RNA Assay Kit in a Qubit 2.0 fluorometer (Thermo Fisher Scientific; catalog # Q32852). The RNA purity was accessed using the NanoPhotometer spectrophotometer (IMPLEN, Westlake Village, CA, USA). The RNA integrity was determined using the RNA Nano 600 Assay Kit on the Bioanalyzer 2100 system (Agilent Technologies, CA, USA). Ribosomal RNA depletion of the samples was performed using Ribo-zero Magnetic Gold Kit (Illumina; catalog # 20037135). Sequencing libraries were generated using NEBNext^®^ Ultra™ II Directional RNA Library Prep Kit for Illumina (New England Biolabs, Ipswich, MA, USA; catalog # E7760L), and were then sequenced using Illumina HiSeq 4000 sequencer for 150 bp read length in paired-end mode (paired-end 150, 2×150). Corresponding raw RNA-sequencing data was generated by the sequencer as FASTQ files for 15 RNA samples (Table 1)^32^.

#### Pre-alignment quality check of dataset

The quality of raw RNA reads in all FASTQ files was evaluated using MultiQC analysis^39^ though the Illumina’s DRAGEN (Dynamic Read Analysis for GENomics) platform (Code Availability 1-2). This evaluation includes per-sequence (**Fig. 2A**) and per-position (**Fig. 2B**) Phred quality score (Q Score) which indicates the accuracy of the raw reads generated by a sequencer^40,41^, the percentage of mapped reads per read group of each sample (**Fig. 2C**) which indicates the quality of the paired-end sequencing^37^, and the total number of sequencing reads obtained per sample (**Fig. 2D**) which reveals an overview of the read depth of the dataset^37^.

**Figure 2.**
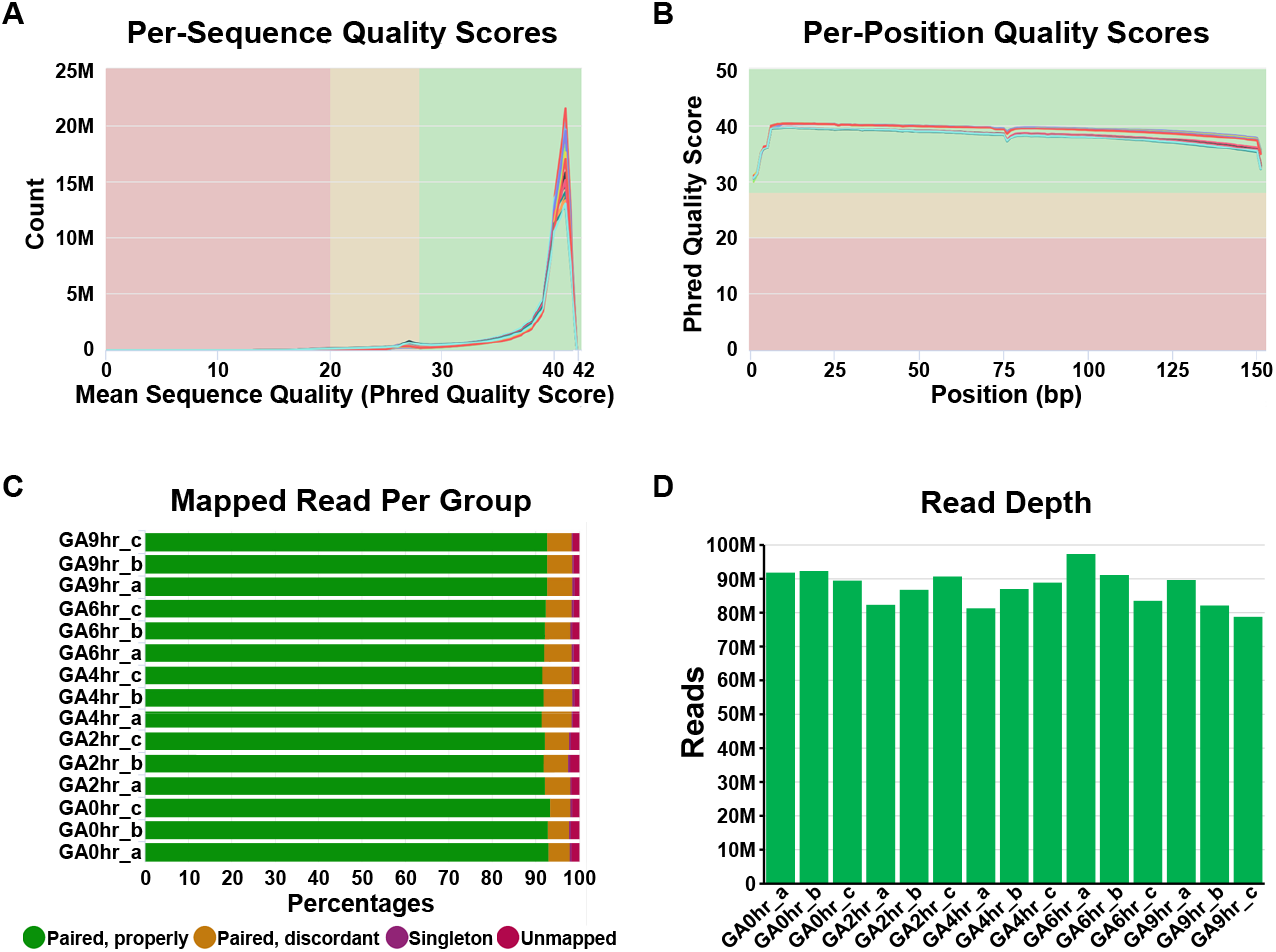
Pre-alignment quality check of dataset. The quality of the raw RNA reads in all FASTQ files of 15 samples in the dataset GSE158788 was evaluated to determine **(A)** mean per-sequence Phred quality scores, **(B)** mean per-position Phred quality scores, **(C)** percentage of paired reads, and **(D)** total number of reads of each sample.

#### Post-alignment quality check of dataset

The post-alignment quality check of the RNA reads in all the FASTQ files was accessed using Partek Flow (Partek Inc., St. Louis, MO, USA) (Code Availability 3) with STAR 2.7.8a aligner package (Code Availability 4) to perform transcript alignment of each sample with the human reference genome hg38^42^. This quality check includes distribution of raw counts of all samples (**Fig. 3A**), coverage breakdown of the counts to reveal the percentage of the reads that fully or partly align with exons or introns (**Fig. 3B**), and sample box plot that illustrates the 10^th^, 25^th^, 50^th^, 75^th^, and 90^th^ percentiles of the counts to show the distribution of normalized gene expression level of each sample (**Fig. 3C**).

**Figure 3.**
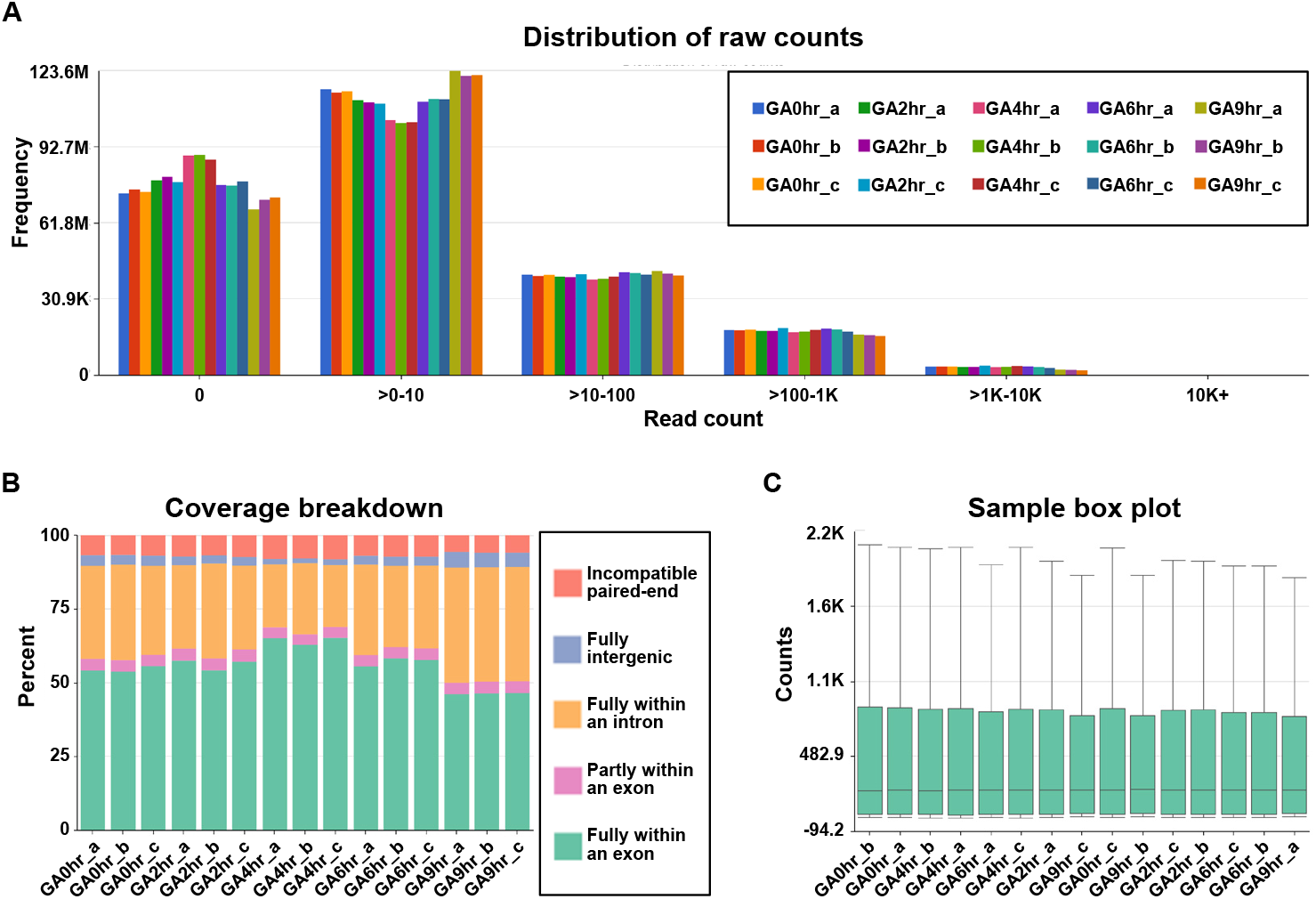
Post-alignment quality check of dataset. The post-alignment quality check of RNA-sequencing data from all 15 samples in the dataset GSE158788 was evaluated to determine **(A)** frequency distribution of raw counts, **(B)** coverage breakdown of the counts, and **(C)** sample box plot that shows 10^th^, 25^th^, 50^th^, 75^th^, and 90^th^ percentiles of the counts after normalization of each sample.

#### Differential gene expression analysis

The characteristics of the dataset and the differential gene expression of coding RNA (messenger RNA, mRNA) and long non-coding RNA (lncRNA) were accessed using Partek Flow (Code Availability 3) with DESeq2 statistical package^43^ (Code Availability 5). These analyses include the three-dimensional scatterplot of principal component analysis (3DPCA)^44,45^ that illustrates the similarity or difference of each sample in the overall dataset with percentage listed on the planes of “X”, “Y”, and “Z” that refer to their corresponding contribution rate (**Fig. 4A**), the hierarchical dendrogram^40,46-48^ that reveals the similarity of gene expression pattens between samples (**Fig.4B**), and differential gene expression of all transcripts (**Fig. 4C**), mRNA (**Fig. 4D**), and lncRNA (**Fig. 4E**) at the 2^nd^ hour (***i*** GA2hr vs GA0hr), 4^th^ hour (***ii*** GA4hr vs GA0hr), 6^th^ hour (***iii*** GA6hr vs GA0hr) and 9^th^ hour (***iv*** GA9hr vs GA0hr) of the gallic acid-induced HeLa cell death process in the dataset.

**Figure 4.**
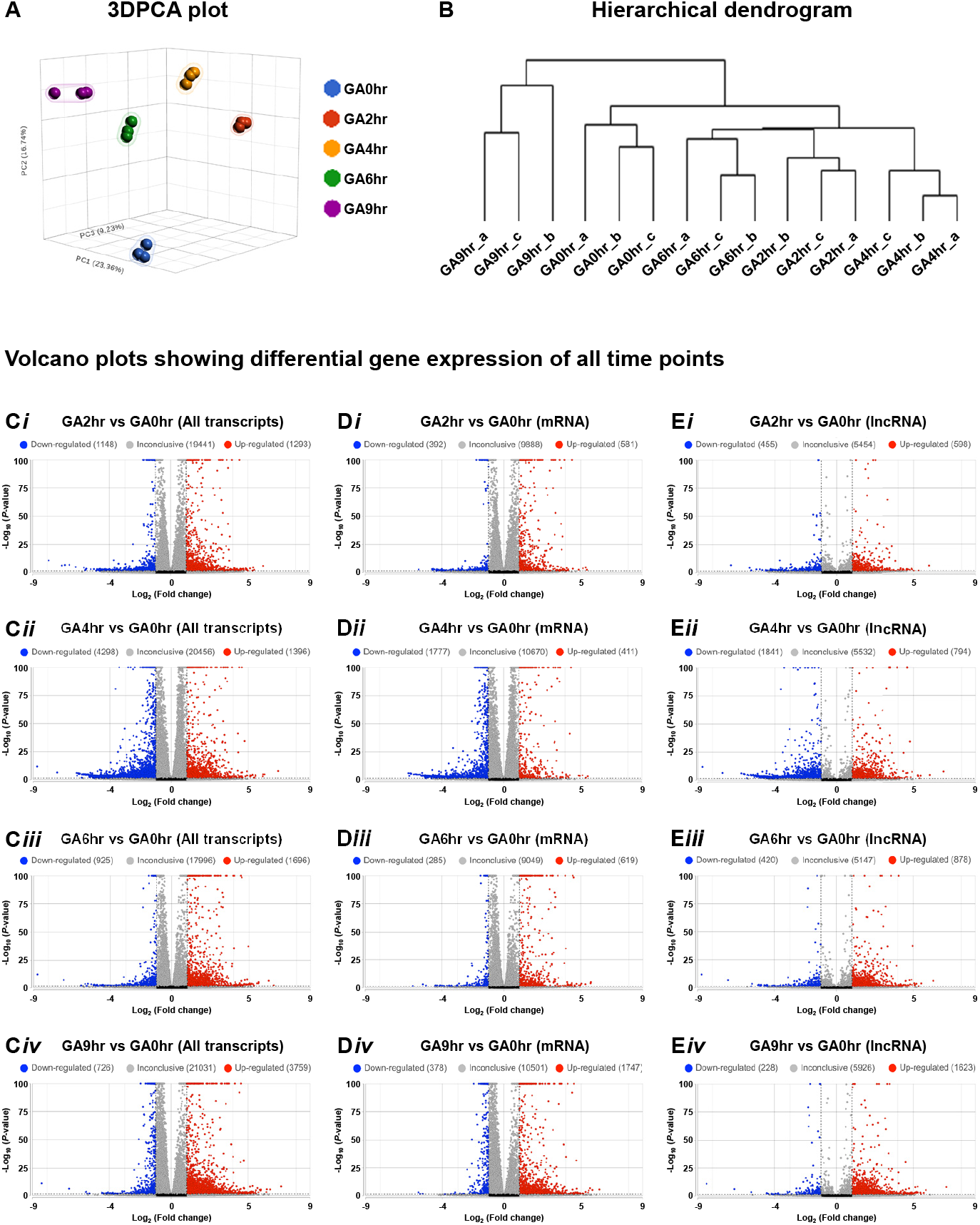
Time-course transcriptomic profiling of gallic acid-induced HeLa cell death. **(A)** Three-dimensional scatterplot of principal component analysis (3DPCA) and **(B)** hierarchical dendrogram of all 15 samples in the dataset GSE158788, and **(C-E)** volcano plots to show the differential gene expression of **(C)** all transcripts, **(D)** mRNA, and **(E)** lncRNA of the 2^nd^ hour (***i*** GA2hr vs GA0hr), 4^th^ hour (***ii*** GA4hr vs GA0hr), 6^th^ hour (***iii*** GA6hr vs GA0hr) and 9^th^ hour (***iv*** GA9hr vs GA0hr) of the gallic acid-induced HeLa cell death process.

## Data Records

All the raw FASTQ files of the dataset for Gene Expression Profile Analysis of Gallic Acid-induced Cell Death Process^32^ was deposited to the Gene Expression Omnibus (GEO) database with the accession number GSE158788^49^.

The dataset is available for public access through this link: https://www.ncbi.nlm.nih.gov/geo/query/acc.cgi?acc=GSE158788 The GEO accession numbers of the raw FASTQ files for all individual samples are listed in Table 1.

### Technical Validation

We performed quality controls on all the 15 samples in our dataset (**Fig. 1** and Table 1, with 5 timepoints, 3 replicates for each timepoint, covering GA0hr, GA2hr, GA4hr, GA6hr, and GA9hr) to evaluate the accuracy of raw RNA-sequencing reads and read depth (**Fig. 2**), distribution and coverage breakdown of raw counts (**Fig. 3**), and sample clustering and differential gene expression analyses (**Fig. 4**) of the dataset^49^.

#### Accuracy of raw RNA sequences

We conducted a pre-alignment quality check of our dataset using MultiQC analysis (**Fig. 2**). The Phred quality score (Q Score) is a standard method widely used to measure the base quality in RNA-sequencing^40,41^. In our dataset, the per-sequence Q Score (**Fig. 2A**) and per-position Q Score (**Fig. 2B**) of most raw reads were significantly higher than Q30 (correct base call by the sequencer is 99.9%, standard from Illumina) and is approaching to Q40 (99.99%). The result of the high correct base call is further validated by the high rate of mapped reads per read group analysis of the paired-end sequencing (**Fig. 2C**), which was higher than 90% for each of the 15 samples.

#### Read depth, and coverage

Our dataset contains at least 78.8 million reads per sample (**Fig. 2D**), which indicates a high read depth that is suitable for studying the molecular mechanisms of cellular process such as differential gene expression^50^. The post-alignment quality check was performed using Partek Flow analysis, which revealed the wide distribution of raw counts (**Fig. 3A**), and the even distribution of the reads that were aligned within exon (e.g., mature RNA) and intron (premature RNA) regions (**Fig. 3B**). Furthermore, the sample box plot illustrates the similar distribution of the counts of all samples in the dataset (**Fig. 3C**). In summary, these validation analyses indicate the high quality of the raw RNA-sequencing dataset in our study.

#### Sample clustering

To illustrate the overall characteristic of the dataset, we evaluated all 15 samples using the three-dimensional scatterplot of principal component analysis (3DPCA) to determine the similarity or difference of each sample in the overall dataset. Our data reveals that all samples from the same time points (biological replicates) closely cluster, while samples belonging to different time points are distinct from others (**Fig. 4A**)^32^. This validation analysis indicates that the gene expression profile of samples at the same time point is similar to each other, and is distinct from that at the different time points. This result is also supported by our hierarchical dendrogram (**Fig. 4B**)^32^ which demonstrates that all the biological replicates are connected with the shortest branch at their corresponding time points, which indicates the highest similarity in their gene expression patterns.

#### Differential gene expression

To validate that our dataset^32^ is suitable for differential gene expression analysis, we applied volcano plots to illustrate the differential expression of all transcripts (**Fig. 4C**) for all four individual time points (GA2hr, GA4hr, GA6hr, GA9hr) for comparison with the control (GA0hr).

Our further validation reveals that the differential expression occurs in both mRNA (**Fig. 4D**) and lncRNA (**Fig. 4E**) at each time point. These analyses demonstrate that the gene expression profile at each stage of gallic acid-induced cell death process is distinct from the control cells that have not been exposed to the gallic acid, thereby indicating that our dataset is suitable for differential gene expression analysis to study the molecular signature of this cell death process.

### Usage Notes

The present study demonstrates that we have generated a high-quality time-course RNA-sequencing dataset revealing the gene expression profile of gallic acid-induced HeLa cell death^32^. Gallic acid is a broad-spectrum anti-cancer compound^1-4^. By comparing the molecular signature of the present dataset generated with human cervical cancer HeLa cells and with those generated in the past and in the future for the cell death response of other human cancer cell types^19-23^ to gallic acid, it will be able to reveal the molecular signature shared by different cancer models, and therefore identify the key regulators and therapeutic targets that mediate gallic acid-induced cancer cell death mechanisms.

Multiple factors could impact the transcriptomic changes that should be taken into consideration for studying the molecular signature of gallic acid. For example, different cancer cell types may respond to gallic acid differently^1-4^. In our present studies, our RNA-Seq detected striking changes in transcription of genes at the early (2^nd^ hour), middle (4^th^, 6^th^ hour), and late (9^th^ hour) stages of cell death in HeLa cells that were exposed to 50μg/mL of gallic acid, in which apoptosis, ferroptosis, and necroptosis were identified^32^. However, differential gene expression was detected in the colon cancer HCT116 cells treated with 50μM of gallic acid for extended time (12^th^ and 24^th^ hour), in which activation of apoptotic cell death pathways was identified^51^. Heterogeneity is the hallmark of cancer cells^25^, and could cause different cellular responses to gallic acid.

Dimethyl sulfoxide (DMSO), a common solvent of gallic acid and numerous agents for biomedical use, could also impact on transcriptomic changes and may cause cytotoxicity. For example, treatment of mouse neuroblastoma N2a cells with 2% DMSO or HeLa cells with 5% DMSO at the final concentration in cell culture medium could affect RNA synthesis^52^ (of note: 0.25% of DMSO was used in our present study). Besides, low concentrations of DMSO (2-4%) could induce cytotoxicity such as in rat retinal ganglion RGC-5 cells *in vitro*, and the high sensitivity of rat retinal cells to DMSO *in vivo* was reported^*53*^. Therefore, proper solvent control is essential for studying the cytotoxicity of gallic acid, and using other solvents for gallic acid such as ethanol could be considered.

The molecular signature of gallic acid-induced cell death can be illustrated by determining the differential gene expression of the datasets using TopHat^54^, Cufflinks^55,56^, HISAT^57^, StringTie^58^, Ballgown^59,60^, and HTSeq^61^. Functional and pathway analyses can be performed using Kyoto Encyclopedia of Genes and Genomes (KEGG)^62,63^, Gene Ontology (GO)^64,65^, Ingenuity Pathway Analysis (IPA)^66^, and WebGestalt^67^. Future analyses of the transcriptomic changes in response gallic acid in different types of cancer cells will reveal the molecular mechanisms governing the anti-cancer property of gallic acid. Therefore, our dataset is a valuable resource with high reuse potential for future studies, and thus facilitate the translation of this natural anti-cancer compound into clinical cancer treatment.

## Code Availability

The pre-alignment quality check, post-alignment quality check, cell clustering, and differential gene expression data analyses were performed using the following software and their default parameters. No custom code was used in the present study.

1) MultiQC, version 1.18: https://multiqc.info/

2) Illumina’s DRAGEN, version 4.2: https://www.illumina.com/products/by-type/informatics-products/dragen-secondary-analysis.html

3) Partek® Flow®, version 10.0: https://www.partek.com/partek-flow/

4) STAR aligner, version 2.7.8a: https://github.com/alexdobin/STAR

5) DESeq2, version 3.18: https://www.bioconductor.org/packages/release/bioc/html/DESeq2.html

## Acknowledgements

This study was supported by the Direct Grant (4053367) of the Chinese University of Hong Kong (CUHK) Research Committee to Dr. Peter Chi Keung Cheung.

## Author contributions

H.M.T. and P.C.K.C. designed the research work, analysed the data, and wrote the manuscript.

H.M.T. performed the experiments.

---

**Table 1.** Gene Expression Omnibus (GEO) accession number of individual samples in the time-course transcriptome dataset for gallic acid-induced HeLa cell death. GEO accession number of the dataset is GSE158788^49^ with 15 samples^68-82^.

## References

1. Subramanian, A.P. et al. Gallic acid: prospects and molecular mechanisms of its anticancer activity. Rsc Advances 5, 35608–35621 (2015).

2. Jiang, Y. et al. Gallic Acid: A Potential Anti-Cancer Agent. Chin J Integr Med 28, 661–671 (2022).

3. Ashrafizadeh, M. et al. Gallic acid for cancer therapy: Molecular mechanisms and boosting efficacy by nanoscopical delivery. Food Chem Toxicol 157, 112576 (2021).

4. Tuli, H.S. et al. Gallic Acid: A Dietary Polyphenol that Exhibits Anti-neoplastic Activities by Modulating Multiple Oncogenic Targets. Anticancer Agents Med Chem 22, 499–514 (2022).

5. Serrano, J., Puupponen-Pimia, R., Dauer, A., Aura, A.M. & Saura-Calixto, F. Tannins: current knowledge of food sources, intake, bioavailability and biological effects. Mol Nutr Food Res 53 Suppl 2, S310–29 (2009).

6. Daglia, M., Di Lorenzo, A., Nabavi, S.F., Talas, Z.S. & Nabavi, S.M. Polyphenols: Well Beyond The Antioxidant Capacity: Gallic Acid and Related Compounds as Neuroprotective Agents: You are What You Eat! Current Pharmaceutical Biotechnology 15, 362–372 (2014).

7. Jara, J.A. et al. Antiproliferative and uncoupling effects of delocalized, lipophilic, cationic gallic acid derivatives on cancer cell lines. Validation in vivo in singenic mice. J Med Chem 57, 2440–54 (2014).

8. Liang, C.Z. et al. Gallic acid induces the apoptosis of human osteosarcoma cells in vitro and in vivo via the regulation of mitogen-activated protein kinase pathways. Cancer Biother Radiopharm 27, 701–10 (2012).

9. Wang, K., Zhu, X., Zhang, K., Zhu, L. & Zhou, F. Investigation of gallic acid induced anticancer effect in human breast carcinoma MCF-7 cells. J Biochem Mol Toxicol 28, 387–93 (2014).

10. Zhao, B. & Hu, M. Gallic acid reduces cell viability, proliferation, invasion and angiogenesis in human cervical cancer cells. Oncol Lett 6, 1749–1755 (2013).

11. Khaledi, H. et al. Antioxidant, cytotoxic activities, and structure-activity relationship of gallic acid-based indole derivatives. Arch Pharm (Weinheim) 344, 703–9 (2011).

12. Ho, H.H. et al. Gallic acid inhibits gastric cancer cells metastasis and invasive growth via increased expression of RhoB, downregulation of AKT/small GTPase signals and inhibition of NF-kappaB activity. Toxicol Appl Pharmacol 266, 76–85 (2013).

13. Inoue, M. et al. Antioxidant, gallic acid, induces apoptosis in HL-60RG cells. Biochem Biophys Res Commun 204, 898–904 (1994).

14. Ohno, Y. et al. Induction of apoptosis by gallic acid in lung cancer cells. Anticancer Drugs 10, 845–51 (1999).

15. Kim, N.S. et al. Gallic acid inhibits cell viability and induces apoptosis in human monocytic cell line U937. J Med Food 14, 240–6 (2011).

16. Varela-Rodriguez, L. et al. Effect of Gallic acid and Myricetin on ovarian cancer models: a possible alternative antitumoral treatment. BMC Complement Med Ther 20, 110 (2020).

17. Kaur, M., Velmurugan, B., Rajamanickam, S., Agarwal, R. & Agarwal, C. Gallic acid, an active constituent of grape seed extract, exhibits anti-proliferative, pro-apoptotic and anti-tumorigenic effects against prostate carcinoma xenograft growth in nude mice. Pharm Res 26, 2133–40 (2009).

18. Ortega, E. et al. Tumoricidal activity of lauryl gallate towards chemically induced skin tumours in mice. Br J Cancer 88, 940–3 (2003).

19. van der Heijden, C.A., Janssen, P.J. & Strik, J.J. Toxicology of gallates: a review and evaluation. Food Chem Toxicol 24, 1067–70 (1986).

20. Rajalakshmi, K., Devaraj, H. & Niranjali Devaraj, S. Assessment of the no-observed-adverse-effect level (NOAEL) of gallic acid in mice. Food Chem Toxicol 39, 919–22 (2001).

21. Niho, N. et al. Subchronic toxicity study of gallic acid by oral administration in F344 rats. Food Chem Toxicol 39, 1063–70 (2001).

22. Vijaya Padma, V., Sowmya, P., Arun Felix, T., Baskaran, R. & Poornima, P. Protective effect of gallic acid against lindane induced toxicity in experimental rats. Food Chem Toxicol 49, 991–8 (2011).

23. Gao, J.Y., Hu, J.X., Hu, D.Y. & Yang, X. A Role of Gallic Acid in Oxidative Damage Diseases: A Comprehensive Review. Natural Product Communications 14(2019).

24. Vasan, N., Baselga, J. & Hyman, D.M. A view on drug resistance in cancer. Nature 575, 299–309 (2019).

25. Hanahan, D. Hallmarks of Cancer: New Dimensions. Cancer Discov 12, 31–46 (2022).

26. Szakacs, G., Paterson, J.K., Ludwig, J.A., Booth-Genthe, C. & Gottesman, M.M. Targeting multidrug resistance in cancer. Nat Rev Drug Discov 5, 219–34 (2006).

27. Scott, E.C. et al. Trends in the approval of cancer therapies by the FDA in the twenty-first century. Nat Rev Drug Discov 22, 625–640 (2023).

28. Carneiro, B.A. & El-Deiry, W.S. Targeting apoptosis in cancer therapy. Nat Rev Clin Oncol 17, 395–417 (2020).

29. Lei, G., Zhuang, L. & Gan, B. Targeting ferroptosis as a vulnerability in cancer. Nat Rev Cancer 22, 381–396 (2022).

30. Gong, Y. et al. The role of necroptosis in cancer biology and therapy. Mol Cancer 18, 100 (2019).

31. Tang, H.M. & Cheung, P.C.K. Gallic Acid Triggers Iron-Dependent Cell Death with Apoptotic, Ferroptotic, and Necroptotic Features. Toxins (Basel) 11(2019).

32. Tang, H.M. & Cheung, P.C.K. Gene expression profile analysis of gallic acid-induced cell death process. Sci Rep 11, 16743 (2021).

33. Vitale, I. et al. Apoptotic cell death in disease-Current understanding of the NCCD 2023. Cell Death Differ 30, 1097–1154 (2023).

34. Stockwell, B.R. Ferroptosis turns 10: Emerging mechanisms, physiological functions, and therapeutic applications. Cell 185, 2401–2421 (2022).

35. Shan, B., Pan, H., Najafov, A. & Yuan, J. Necroptosis in development and diseases. Genes Dev 32, 327–340 (2018).

36. Masters, J.R. HeLa cells 50 years on: the good, the bad and the ugly. Nat Rev Cancer 2, 315–9 (2002).

37. Stark, R., Grzelak, M. & Hadfield, J. RNA sequencing: the teenage years. Nat Rev Genet 20, 631–656 (2019).

38. Mattick, J.S. et al. Long non-coding RNAs: definitions, functions, challenges and recommendations. Nat Rev Mol Cell Biol 24, 430–447 (2023).

39. Ewels, P., Magnusson, M., Lundin, S. & Kaller, M. MultiQC: summarize analysis results for multiple tools and samples in a single report. Bioinformatics 32, 3047–8 (2016).

40. Li, W.V. & Li, J.J. Modeling and analysis of RNA-seq data: a review from a statistical perspective. Quant Biol 6, 195–209 (2018).

41. Richterich, P. Estimation of errors in “raw” DNA sequences: a validation study. Genome Res 8, 251–9 (1998).

42. Genome assembly GRCh38.p14. National Library of Medicine, https://www.ncbi.nlm.nih.gov/datasets/genome/GCF_000001405.40/ (2022).

43. Love, M.I., Huber, W. & Anders, S. Moderated estimation of fold change and dispersion for RNA-seq data with DESeq2. Genome Biol 15, 550 (2014).

44. Ringner, M. What is principal component analysis? Nat Biotechnol 26, 303–4 (2008).

45. Jolliffe, I.T. & Cadima, J. Principal component analysis: a review and recent developments. Philos Trans A Math Phys Eng Sci 374, 20150202 (2016).

46. Conesa, A. et al. A survey of best practices for RNA-seq data analysis. Genome Biol 17, 13 (2016).

47. Van den Berge, K. et al. RNA Sequencing Data: Hitchhiker’s Guide to Expression Analysis. Annual Review of Biomedical Data Science, Vol 2, 2019 2, 139–173 (2019).

48. Boutros, P.C. & Okey, A.B. Unsupervised pattern recognition: an introduction to the whys and wherefores of clustering microarray data. Brief Bioinform 6, 331–43 (2005).

49. Tang, H.M. & Cheung, P.C.K. Gene Expression Profile Analysis of Gallic Acid-induced Cell Death Process. Gene Expression Omnibus, https://www.ncbi.nlm.nih.gov/geo/query/acc.cgi?acc=GSE158788 (2021).

50. Sims, D., Sudbery, I., Ilott, N.E., Heger, A. & Ponting, C.P. Sequencing depth and coverage: key considerations in genomic analyses. Nat Rev Genet 15, 121–32 (2014).

51. Yang, C. et al. Transcriptome analysis reveals GA induced apoptosis in HCT116 human colon cancer cells through calcium and p53 signal pathways. RSC Adv 8, 12449–12458 (2018).

52. Bolduc, L., Labrecque, B., Cordeau, M., Blanchette, M. & Chabot, B. Dimethyl sulfoxide affects the selection of splice sites. J Biol Chem 276, 17597–602 (2001).

53. Galvao, J. et al. Unexpected low-dose toxicity of the universal solvent DMSO. FASEB J 28, 1317–30 (2014).

54. Trapnell, C., Pachter, L. & Salzberg, S.L. TopHat: discovering splice junctions with RNA-Seq. Bioinformatics 25, 1105–11 (2009).

55. Trapnell, C. et al. Transcript assembly and quantification by RNA-Seq reveals unannotated transcripts and isoform switching during cell differentiation. Nat Biotechnol 28, 511–5 (2010).

56. Trapnell, C. et al. Differential gene and transcript expression analysis of RNA-seq experiments with TopHat and Cufflinks. Nat Protoc 7, 562–78 (2012).

57. Kim, D., Langmead, B. & Salzberg, S.L. HISAT: a fast spliced aligner with low memory requirements. Nat Methods 12, 357–60 (2015).

58. Pertea, M. et al. StringTie enables improved reconstruction of a transcriptome from RNA-seq reads. Nat Biotechnol 33, 290–5 (2015).

59. Frazee, A.C. et al. Ballgown bridges the gap between transcriptome assembly and expression analysis. Nat Biotechnol 33, 243–6 (2015).

60. Pertea, M., Kim, D., Pertea, G.M., Leek, J.T. & Salzberg, S.L. Transcript-level expression analysis of RNA-seq experiments with HISAT, StringTie and Ballgown. Nat Protoc 11, 1650–67 (2016).

61. Anders, S., Pyl, P.T. & Huber, W. HTSeq--a Python framework to work with high-throughput sequencing data. Bioinformatics 31, 166–9 (2015).

62. Kanehisa, M. & Goto, S. KEGG: kyoto encyclopedia of genes and genomes. Nucleic Acids Res 28, 27–30 (2000).

63. Kanehisa, M., Furumichi, M., Tanabe, M., Sato, Y. & Morishima, K. KEGG: new perspectives on genomes, pathways, diseases and drugs. Nucleic Acids Res 45, D353–D361 (2017).

64. Ashburner, M. et al. Gene ontology: tool for the unification of biology. The Gene Ontology Consortium. Nat Genet 25, 25–9 (2000).

65. Gene Ontology, C. et al. The Gene Ontology knowledgebase in 2023. Genetics 224(2023).

66. Kramer, A., Green, J., Pollard, J., Jr. & Tugendreich, S. Causal analysis approaches in Ingenuity Pathway Analysis. Bioinformatics 30, 523–30 (2014).

67. Liao, Y., Wang, J., Jaehnig, E.J., Shi, Z. & Zhang, B. WebGestalt 2019: gene set analysis toolkit with revamped UIs and APIs. Nucleic Acids Res 47, W199–W205 (2019).

68. Tang, H. M. & Cheung, P.C.K. Gene Expression Omnibus. https://identifiers.org/geo:GSM4810497 (2021).

69. Tang, H. M. & Cheung, P.C.K. Gene Expression Omnibus. https://identifiers.org/geo:GSM4810498 (2021).

70. Tang, H. M. & Cheung, P.C.K. Gene Expression Omnibus. https://identifiers.org/geo:GSM4810499 (2021).

71. Tang, H. M. & Cheung, P.C.K. Gene Expression Omnibus. https://identifiers.org/geo:GSM4810500 (2021).

72. Tang, H. M. & Cheung, P.C.K. Gene Expression Omnibus. https://identifiers.org/geo:GSM4810501 (2021).

73. Tang, H. M. & Cheung, P.C.K. Gene Expression Omnibus. https://identifiers.org/geo:GSM4810502 (2021).

74. Tang, H. M. & Cheung, P.C.K. Gene Expression Omnibus. https://identifiers.org/geo:GSM4810503 (2021).

75. Tang, H. M. & Cheung, P.C.K. Gene Expression Omnibus. https://identifiers.org/geo:GSM4810504 (2021).

76. Tang, H. M. & Cheung, P.C.K. Gene Expression Omnibus. https://identifiers.org/geo:GSM4810505 (2021).

77. Tang, H. M. & Cheung, P.C.K. Gene Expression Omnibus. https://identifiers.org/geo:GSM4810506 (2021).

78. Tang, H. M. & Cheung, P.C.K. Gene Expression Omnibus. https://identifiers.org/geo:GSM4810507 (2021).

79. Tang, H. M. & Cheung, P.C.K. Gene Expression Omnibus. https://identifiers.org/geo:GSM4810508 (2021).

80. Tang, H. M. & Cheung, P.C.K. Gene Expression Omnibus. https://identifiers.org/geo:GSM4810509 (2021).

81. Tang, H. M. & Cheung, P.C.K. Gene Expression Omnibus. https://identifiers.org/geo:GSM4810510 (2021).

82. Tang, H. M. & Cheung, P.C.K. Gene Expression Omnibus. https://identifiers.org/geo:GSM4810511 (2021).

